# Inverse regulation of *Vibrio cholerae* biofilm dispersal by polyamine signals

**DOI:** 10.1101/2020.12.06.413526

**Authors:** Andrew A. Bridges, Bonnie L. Bassler

## Abstract

The global pathogen *Vibrio cholerae* undergoes cycles of biofilm formation and dispersal in the environment and the human host. Little is understood about biofilm dispersal. Here, we show that MbaA, a periplasmic polyamine sensor, and PotD1, a polyamine importer, regulate *V. cholerae* biofilm dispersal. Spermidine, a commonly produced polyamine, drives *V. cholerae* dispersal, whereas norspermidine, an uncommon polyamine produced by vibrios, inhibits dispersal. Spermidine and norspermidine differ by one methylene group. Both polyamines function to control dispersal via periplasmic detection by MbaA and subsequent signal relay. Biofilm dispersal fails in the absence of PotD1 because reuptake of endogenously produced norspermidine does not occur, so it accumulates in the periplasm where it stimulates MbaA. These results suggest that *V. cholerae* uses MbaA to monitor environmental polyamines, blends of which potentially provide information about numbers of ‘self’ and ‘other’. This information is used to dictate whether or not to disperse from biofilms.

## Introduction

Bacteria frequently colonize environmental habitats and infection sites by forming surface-attached multicellular communities called biofilms (Flemming et al., 2016). Participating in the biofilm lifestyle allows bacteria to collectively acquire nutrients and resist threats (Flemming et al., 2016). By contrast, the individual free-swimming state allows bacteria to roam. The global pathogen *Vibrio cholerae* undergoes repeated rounds of biofilm formation and disassembly, and both biofilm formation and biofilm exit are central to disease transmission as *V. cholerae* alternates between the marine niche and the human host (Conner et al., 2016; Gallego-Hernandez et al., 2020; Tamayo et al., 2010). The *V. cholerae* biofilm lifecycle occurs in three stages. First, a founder cell attaches to a substate, second, through cell division and extracellular matrix secretion, the biofilm matures, and finally, when the appropriate environmental conditions are detected, cells leave the biofilm through a process termed dispersal (Bridges et al., 2020). Bacterial cells liberated through dispersal can depart their current environment and repeat the lifecycle in a new niche, such as in a new host (Guilhen et al., 2017). In general, the initial stages of the biofilm lifecycle, from attachment through maturation, have been well studied in *V. cholerae* and other bacterial species, whereas the dispersal phase remains underexplored.

To uncover the mechanisms controlling *V. cholerae* biofilm dispersal, we recently developed a brightfield microscopy assay that allows us to follow the entire biofilm lifecycle (Bridges and Bassler, 2019). To identify the components that orchestrate *V. cholerae* biofilm dispersal, we combined this assay with mutagenesis and high-content imaging to identify mutants that failed to properly disperse (Bridges et al., 2020). Our screen revealed genes encoding proteins that fall into three functional groups: signal transduction, matrix degradation, and cell motility. Our hypothesis is that these classes of components act in sequence to drive biofilm dispersal. First, the cues that induce dispersal are detected by the signal transduction proteins. Second, activation of matrix digestion components occurs. Finally, cell motility engages and permits cells to swim away from the disassembling biofilm. The majority of the genes identified in the screen encoded signal transduction proteins. Elsewhere, we characterized one of the signaling systems identified in the screen, a new two-component phospho-relay that we named DbfS-DbfR (for Dispersal of Biofilm Sensor and Dispersal of Biofilm Regulator) that controls biofilm dispersal (Bridges et al., 2020). Of the remaining signaling proteins identified in the screen, four are proteins involved in regulating production/degradation of, or responding to, the second messenger molecule cyclic diguanylate (c-di-GMP).

C-di-GMP has a known role in the regulation of biofilm formation in many bacteria including *V. cholerae* in which low c-di-GMP levels correlate with motility and high c-di-GMP levels promote surface attachment and matrix production, and thus biofilm formation (Conner et al., 2017). C-di-GMP is synthesized by enzymes that contain catalytic GGDEF domains that have diguanylate cyclase activity (Conner et al., 2017). C-di-GMP is degraded by proteins with EAL or HD-GYP domains that possess phosphodiesterase activity (Conner et al., 2017). *V. cholerae* encodes >50 proteins harboring one or more of these domains, underscoring the global nature of c-di-GMP signaling in this pathogen. The activities of these enzymes are often regulated by environmental stimuli through ligand binding sensory domains, however, in most cases, the identities of ligands for GGDEF- and EAL-containing enzymes are unknown (Römling et al., 2013). Of the c-di-GMP regulatory proteins identified in our dispersal screen only one, the hybrid GGDEF/EAL protein, MbaA, which has previously been shown to regulate *V. cholerae* biofilm formation (Bomchil et al., 2003), has a defined environmental stimulus: MbaA responds to the ubiquitous family of small molecules called polyamines, which are involved in biological processes ranging from signal transduction to protein synthesis to siderophore production (Karatan et al., 2005; McGinnis et al., 2009; Michael, 2018). Our dispersal screen also identified PotD1, a periplasmic binding protein that mediates polyamine import and has also been shown to control *V. cholerae* biofilm formation (Cockerell et al., 2014; McGinnis et al., 2009). Thus, identification of these two genes in our screen suggested that beyond roles in biofilm formation, polyamine detection could be central to biofilm dispersal. In the present work, we explore this new possibility.

Previous work showed that norspermidine, a rare polyamine produced by Vibrionaceae and select other organisms, promotes biofilm formation while the nearly ubiquitously-made polyamine spermidine, which has been reported not to be produced by *V. cholerae*, represses biofilm formation (Fig. 1) (Karatan et al., 2005; McGinnis et al., 2009). The hypothesis is that *V. cholerae* detects norspermidine as a measure of ‘self’ and spermidine as a measure of ‘other’ to assess the species composition of the vicinal community and it funnels that information internally to determine whether or not to make a biofilm (Wotanis et al., 2017). As mentioned above, information contained in extracellular polyamines is transduced internally by MbaA, which is embedded in the inner membrane. MbaA contains a periplasmic domain that is thought to interact with the periplasmic binding protein NpsS (Cockerell et al., 2014; Karatan et al., 2005). MbaA also possesses cytoplasmic GGDEF (SGDEF in MbaA) and EAL (EVL in MbaA) domains (Fig. 1). Genetic evidence suggests that when NpsS is bound to norspermidine, the complex associates with MbaA and biofilm formation is promoted (Cockerell et al., 2014). The presumption is that NpsS interaction with MbaA inhibits MbaA phosphodiesterase activity, leading to an increase in c-di-GMP levels, and in turn, elevated biofilm formation (Fig. 1, Left) (Cockerell et al., 2014). Apo-NpsS or NpsS bound to spermidine does not bind to MbaA, and thus, under this condition, biofilm formation is reduced (Cockerell et al., 2014). The hypothesis in this case is that MbaA phosphodiesterase activity is uninhibited, which leads to decreased c-di-GMP levels and repression of biofilm formation (Fig. 1, Right). Currently, it is not known whether or not MbaA has diguanylate cyclase activity. The purified MbaA cytoplasmic domain functions as a phosphodiesterase *in vitro* (Cockerell et al., 2014).

**Fig. 1.**
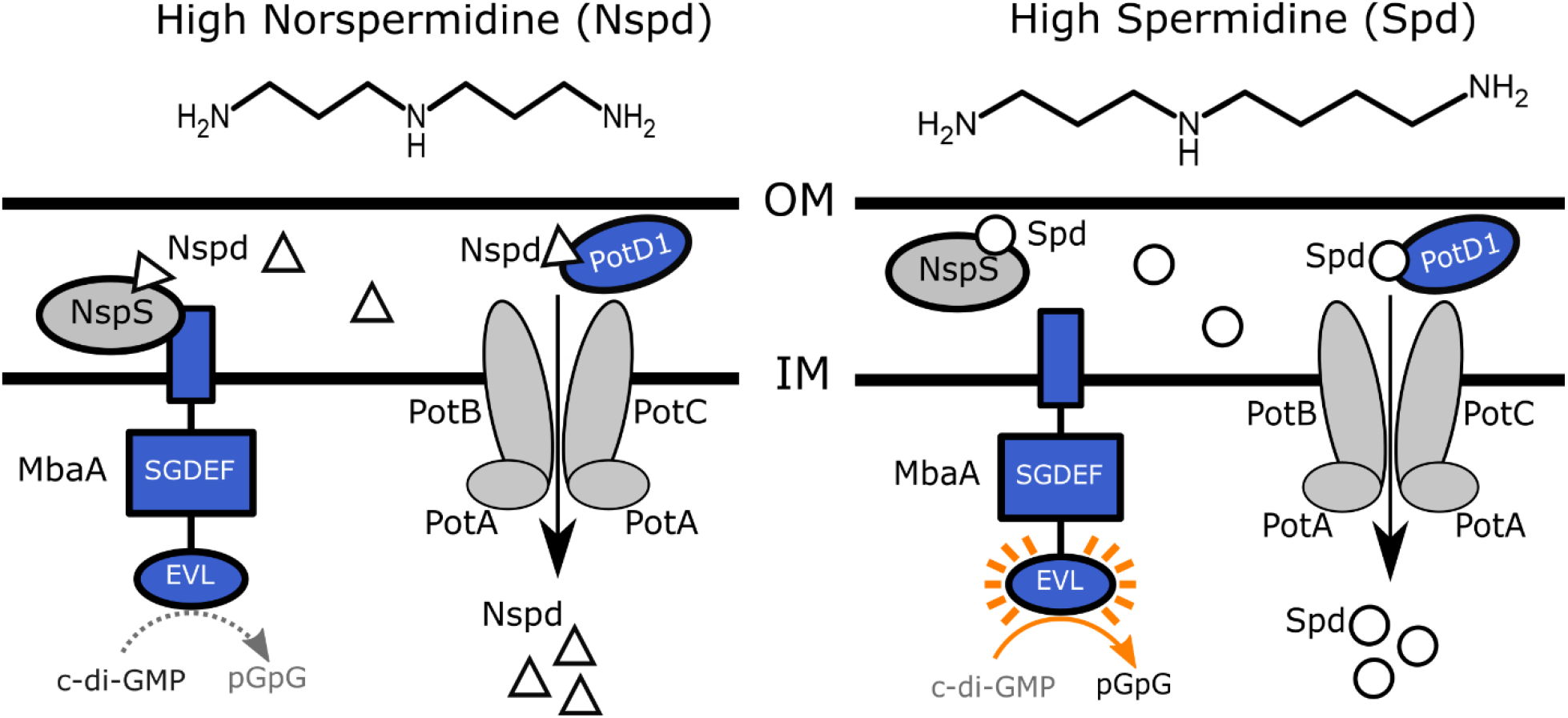
Polyamine sensing in *V. cholerae*. Schematic showing the previously proposed polyamine detection and import mechanisms in *V. cholerae*. Norspermidine (Nspd, triangles) promotes biofilm formation and spermidine (Spd, circles) represses biofilm formation. See text for details. OM = outer membrane; IM = inner membrane.

In a proposed second regulatory mechanism, both polyamines are thought to control biofilm formation via import through the inner membrane ABC transporter, PotABCD1 (McGinnis et al., 2009). Following internalization, via an undefined cytoplasmic mechanism, polyamines are proposed to modulate biofilm formation (Fig. 1) (McGinnis et al., 2009). This hypothesis was based on the finding that elimination of PotD1 that is required for import of norspermidine and spermidine resulted in elevated biofilm formation. Importantly, the studies reporting the MbaA and PotD1 findings used quorum-sensing deficient *V. cholerae* strains (Joelsson et al., 2006) that likely cannot disperse from biofilms and end-point biofilm assays and so were unable to differentiate between enhanced biofilm formation and the failure to disperse.

Here, we combine real-time biofilm lifecycle measurements, mutagenesis, and a reporter of cytoplasmic c-di-GMP levels to define the roles polyamines play over the entire biofilm lifecycle. In wildtype *V. cholerae*, exogenous norspermidine and spermidine inversely alter cytoplasmic c-di-GMP levels and they exert their effects through the NspS-MbaA circuit. Norspermidine promotes biofilm formation and suppresses biofilm dispersal. Spermidine represses biofilm formation and promotes biofilm dispersal. Both the MbaA SGDEF and EVL domains are required for polyamine control of the biofilm lifecycle. We also show that MbaA synthesizes c-di-GMP in the presence of norspermidine and degrades c-di-GMP in the presence of spermidine and that is what drives *V. cholerae* to form and disperse from biofilms, respectively. When MbaA is absent, *V. cholerae* is unable to alter the biofilm lifecycle in response to extracellular polyamines. We demonstrate that polyamine internalization via PotD1 is not required to promote *V. cholerae* entrance or exit from biofilms, but rather, periplasmic detection of polyamines by MbaA is the key regulatory step. Specifically, the Δ*potD1* mutant fails to disperse because it is unable to reuptake self-secreted norspermidine. The consequence is that excess periplasmic norspermidine accumulates, is detected by MbaA, and leads to production of c-di-GMP and suppression of biofilm dispersal. Collectively, our work reveals the mechanisms by which *V. cholerae* detects and transduces polyamine signals to modulate its biofilm lifecycle. We propose that the polyamine sensing mechanisms revealed in this study allow *V. cholerae* to distinguish relatives from competitors and potentially the presence of predators in the vicinity and, in response, modify its biofilm lifecycle to appropriately colonize territory or disperse from an existing community.

## Results

### Polyamine signaling proteins MbaA and PotD1 regulate biofilm dispersal through c-di-GMP

Our combined mutagenesis-imaging screen identified the inner membrane polyamine sensor, MbaA, and the periplasmic polyamine binding protein PotD1 as essential for proper *V. cholerae* biofilm dispersal motivating us to explore the mechanisms underlying these effects. To probe polyamine signaling across the full biofilm lifecycle, we used our established brightfield imaging assay. In the case of WT *V. cholerae*, peak biofilm biomass is reached at ∼8-9 h of growth, and subsequently, dispersal occurs, and is completed by ∼12 h post inoculation (Fig. 2A, B). Deletion of *mbaA* caused a mild biofilm dispersal defect with no detectable difference in peak biofilm biomass compared to WT, a less than 1 h delay in the onset of dispersal, and 27% biomass remaining at 16 h (Fig. 2A, B). Thus, MbaA regulates the dispersal phase of the biofilm lifecycle. The Δ*potD1* mutant exhibited a 60% greater peak biofilm biomass than WT and nearly all of the biomass remained at 16 h, indicating that PotD1 both represses biofilm formation and promotes biofilm dispersal (Fig. 2A, B). The differences in severity between the dispersal phenotypes of the Δ*mbaA* and Δ*potD1* mutants are noteworthy because these strains behaved similarly in the previous end-point biofilm formation assays (McGinnis et al., 2009).

**Fig. 2.**
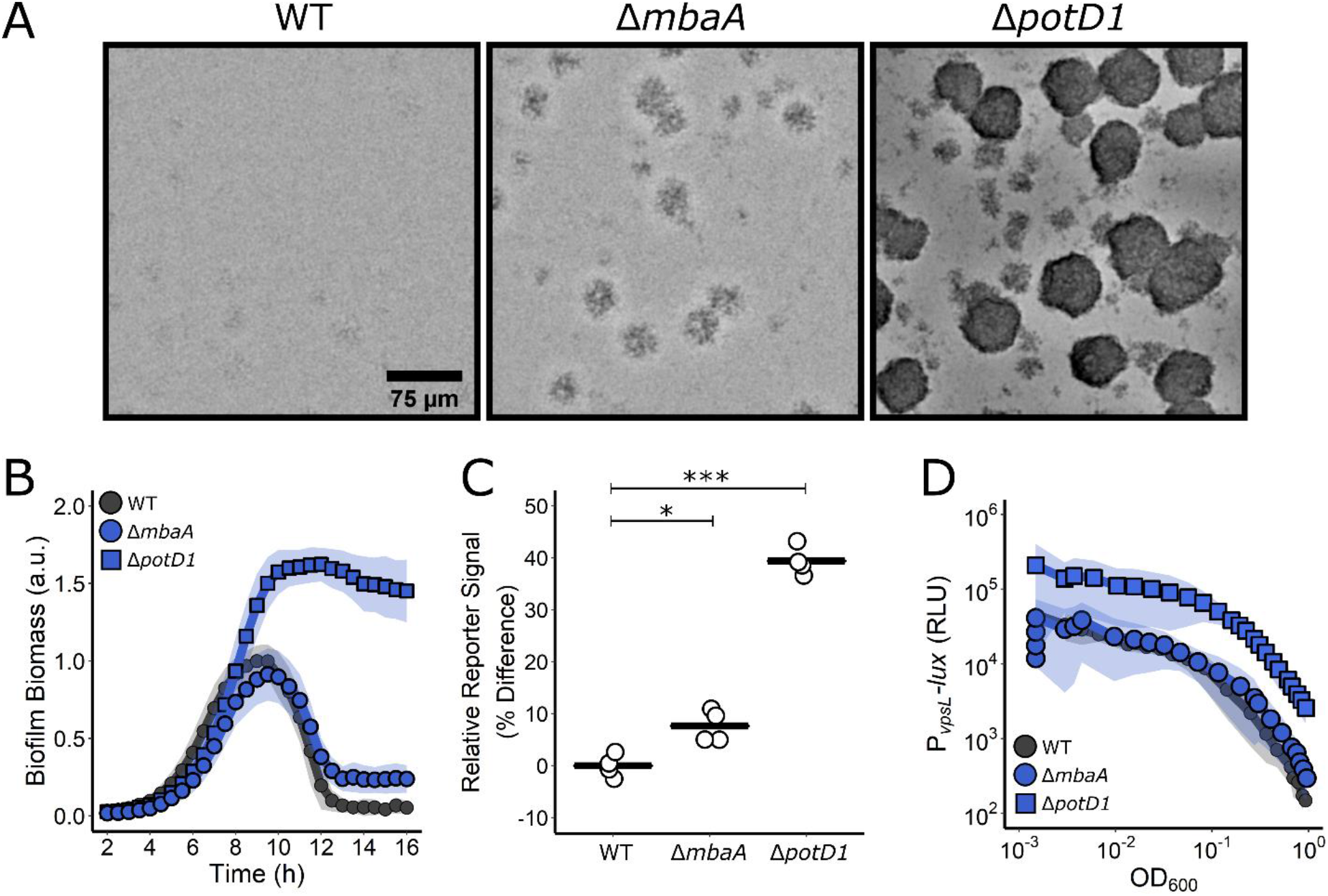
Polyamine signaling regulates *V. cholerae* biofilm dispersal. (A) Representative images of the designated *V. cholerae* strains at 16 h. (B) Quantitation of biofilm biomass over time measured by time-lapse microscopy for WT *V. cholerae* and the designated mutants. In all cases, *N* = 3 biological and *N* = 3 technical replicates, ± SD (shaded). a.u., arbitrary unit. (C) Relative c-di-GMP reporter signal for the indicated strains. Values are expressed as the percentage difference relative to the WT strain. *N* = 4 biological replicates. The black bar shows the sample mean. Unpaired t-tests were performed for statistical analysis, with P values denoted as *P < 0.05; **P < 0.005; *** P < 0.0005. (D) The corresponding *PvpsL-lux* output for strains and growth conditions as in C. For *vpsL-lux* measurements, *N* = 3 biological replicates, ± SD (shaded). RLU, relative light units.

MbaA has previously been shown possess phosphodiesterase activity *in vitro* (Cockerell et al., 2014), thus, we reasoned that MbaA functions by altering cytoplasmic c-di-GMP levels, the consequence of which is changes in expression of genes encoding biofilm components. As far as we are aware, no connection has yet been made between PotD1 and cytoplasmic c-di-GMP levels. To determine if the observed biofilm dispersal defects in the Δ*mbaA* and Δ*potD1* mutants track with altered c-di-GMP levels, we compared the relative cytoplasmic c-di-GMP levels in the WT, Δ*mbaA*, and Δ*potD1* strains. To do this, we employed a fluorescent reporter construct in which expression of *turboRFP* is controlled by two c-di-GMP-responsive riboswitches (Zhou et al., 2016). The TurboRFP signal is normalized to constitutively produced amCyan encoded on the same plasmid. Previous studies have demonstrated a linear relationship between reporter output and c-di-GMP levels measured by mass spectrometry (Zhou et al., 2016). Indeed, the reporter showed that deletion of *mbaA* caused only a moderate increase in cytoplasmic c-di-GMP (8% higher than WT), while the Δ*potD1* mutant produced 39% more signal than WT (Fig. 2C). Thus, MbaA, as expected, mediates changes in c-di-GMP levels, and moreover PotD1-mediated import of polyamines also influences cytoplasmic c-di-GMP concentrations.

In *V. cholerae*, increased cytoplasmic c-di-GMP levels are associated with elevated extracellular matrix production (called VPS for vibrio polysaccharide) and, in turn, increased biofilm formation. Using a *P*_*vpsL*_*-lux* promoter fusion that reports on the major matrix biosynthetic operon, we previously showed that matrix production decreases as cells transition from the biofilm to the planktonic state, suggesting that repression of matrix production correlates with biofilm dispersal. We wondered how the increased c-di-GMP levels present in the Δ*mbaA* and Δ*potD1* mutants impinged on *vpsL* expression. Using the *vpsL-lux* reporter, we found that the light production patterns mirrored the severities of the dispersal phenotypes and the magnitudes of changes in cytoplasmic c-di-GMP levels: the Δ*mbaA* mutant had a light production profile similar to WT, while the Δ*potD1* mutant produced 10-fold more light than WT throughout growth. These results show that the Δ*mbaA* mutant makes normal levels and the Δ*potD1* mutant produces excess matrix (Fig. 2D).

As mentioned in the Introduction, our screen revealed three additional genes in c-di-GMP signaling pathways involved in *V. cholerae* biofilm dispersal. While not the focus of the current work, we performed preliminary characterization by deleting these genes and making measurements of their biofilm lifecycle phenotypes, assessing them for changes in cytoplasmic c-di-GMP levels, and quantifying their matrix production profiles (Fig. S1A-D).

### Norspermidine and spermidine inversely regulate *V. cholerae* biofilm dispersal

Above we show that polyamine signaling proteins are involved in driving *V. cholerae* biofilm dispersal. Previous studies demonstrated that norspermidine and spermidine have opposing effects on biofilm formation in end-point assays (Karatan et al., 2005; McGinnis et al., 2009). We wondered if and how these two polyamines affect each stage of the biofilm lifecycle – biofilm formation *and* biofilm dispersal. To investigate their roles, we assayed the biofilm lifecycle in WT *V. cholerae* and our mutants following exogenous administration of norspermidine and spermidine, alone and in combination. Addition of 100 µM norspermidine strongly promoted biofilm formation and completely prevented biofilm dispersal while 100 µM spermidine dramatically reduced biofilm formation and promoted premature biofilm dispersal (Fig. 3A, Video 1). *vpsL-lux* expression increased by > 10-fold following norspermidine treatment and decreased >10-fold following spermidine treatment (Fig. 3B). These results show that norspermidine and spermidine have opposing activities with regard to biofilm formation and dispersal: norspermidine drives biofilm formation by inducing matrix production and this prevents biofilm dispersal and spermidine, by suppressing matrix production prevents biofilm formation and drives biofilm dispersal.

**Fig. 3.**
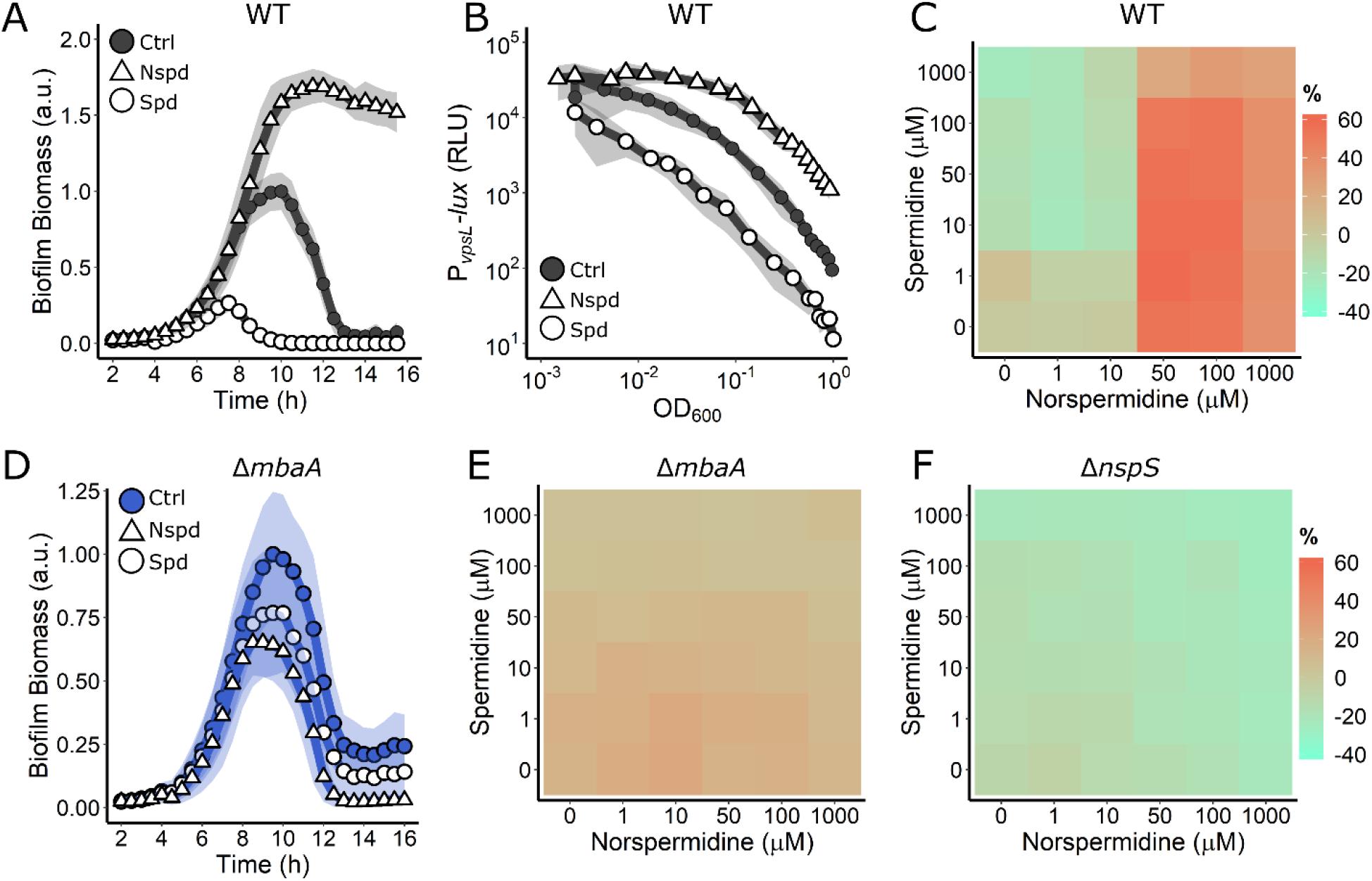
Periplasmic detection of polyamines controls *V. cholerae* biofilm dispersal. (A) Quantitation of biofilm biomass over time measured by time-lapse microscopy following addition of water (Ctrl), 100 µM norspermidine, or 100 µM spermidine to WT *V. cholerae*. (B) Light output from the *PvpsL-lux* reporter for the treatments in A over the growth curve. (C) C-di-GMP reporter output at the indicated polyamine concentrations for WT *V. cholerae*. Relative reporter signal (% difference) is displayed as a heatmap (teal and orange represent the lowest and highest reporter output, respectively). (D) As in A for the Δ*mbaA* mutant. (E) As in C for the Δ*mbaA* mutant. (F) As in C for the Δ*nspS* mutant. Biofilm biomass data are represented as means normalized to the peak biofilm biomass of the Ctrl condition. In all biofilm biomass measurements, *N* = 3 biological and *N* = 3 technical replicates, ± SD (shaded). a.u., arbitrary unit. In *vpsL-lux* measurements, *N* = 3 biological replicates, ± SD (shaded). RLU, relative light units. For the c-di-GMP reporter assays, values are expressed as the percentage difference relative to the untreated WT strain, allowing comparisons to be made across all heatmaps in all figures in this manuscript. The same color bar applies to all heatmaps in this manuscript. For each condition, *N* = 3 biological replicates. Numerical values and associated standard deviations are available in Table S1.

We wondered by what mechanism the information encoded in exogenous polyamines was translated into alterations in matrix gene expression and subsequent changes in biofilm dispersal. We reasoned that MbaA and/or PotD1-mediated changes in cytoplasmic c-di-GMP levels could be responsible. To test this possibility, we assessed how changes in cytoplasmic c-di-GMP levels tracked with changes in extracellular polyamine levels by measuring the c-di-GMP reporter output in response to supplied mixtures of norspermidine and spermidine (Fig. 3C). In the heatmaps, teal and orange represent the lowest and highest values of reporter output, respectively. When provided alone and above a concentration of 10 µM, norspermidine and spermidine strongly increased and decreased, respectively, c-di-GMP reporter activity. At the limit, relative to the WT, norspermidine increased the c-di-GMP reporter output by ∼60% while spermidine reduced reporter output by ∼25%. Notably, when norspermidine was present above 50 µM, it overrode the effect of exogenous addition of spermidine, irrespective of spermidine concentration (Fig. 3C). Thus, *V. cholerae* cytoplasmic c-di-GMP levels are responsive to extracellular blends of polyamines. When norspermidine is abundant, *V. cholerae* produces high levels of c-di-GMP, matrix expression is increased, and biofilms and do not disperse. When spermidine is abundant and norspermidine is absent or present below a threshold concentration, *V. cholerae* c-di-GMP levels drop, matrix production is repressed, and biofilm dispersal occurs.

### MbaA transduces external polyamine information to control biofilm dispersal

Our data show that both the MbaA periplasmic polyamine sensor and the PotD1 polyamine importer are required for biofilm dispersal. Our next goal was to distinguish the contribution of periplasmic detection from cytoplasmic import of polyamines to the biofilm lifecycle. We began with periplasmic polyamine detection, mediated by NpsS together with MbaA. We supplied norspermidine or spermidine to the Δ*mbaA* mutant and monitored biofilm biomass over time. To our surprise, the Δ*mbaA* mutant was impervious to the addition of either polyamine as both polyamines caused only a very modest reduction in overall biofilm biomass, and dispersal timing resembled the untreated Δ*mbaA* control (Fig. 3D, Video 1). Consistent with this result, titration of the polyamines alone and in combination onto the Δ*mbaA* strain carrying the c-di-GMP reporter did not substantially alter reporter output (Fig. 3E). These results show that the dynamic response of WT to polyamines requires MbaA. We suspect that the minor reduction in biofilm production that occurred in the Δ*mbaA* mutant when supplied polyamines is due to non-specific effects, perhaps via interaction of polyamines with the negatively charged VPS. We next investigated the role of the polyamine periplasmic binding protein NspS that transmits polyamine information to MbaA. In the absence of its partner polyamine binding protein, NspS, MbaA is thought to function as a constitutive phosphodiesterase (Cockerell et al., 2014). Consistent with this model, deletion of *nspS* in an otherwise WT strain reduced overall peak biofilm biomass by 65% and dispersal initiated 4 h prior to when dispersal begins in the WT (Fig. S2). In the Δ*nspS* mutant, c-di-GMP levels were lower than in the WT as judged by the c-di-GMP reporter (Fig. 3F) showing that MbaA is locked as a constitutive phosphodiesterase. Exogenous addition of polyamines had no effect on c-di-GMP levels (Fig. 3F). Together, these findings demonstrate that the WT *V. cholerae* response to polyamines is controlled by the NpsS-MbaA polyamine sensing circuit.

### Both the MbaA EVL and SGDEF domains are required for MbaA to detect polyamines

We wondered how the putative MbaA phosphodiesterase and diguanylate cyclase enzymatic activities contribute to the *V. cholerae* responses to norspermidine and spermidine. The cytoplasmic domain of MbaA exhibits phosphodiesterase but not diguanylate cyclase activity *in vitro* (Cockerell et al., 2014). In the presence of an intact regulatory domain. We reasoned that it is possible that MbaA possesses both enzymatic activities, with the activity of each domain inversely controlled by NpsS binding. To probe the role of each domain, we introduced inactivating point mutations in the catalytic motifs. To ensure that our mutations did not destabilize MbaA, we first fused MbaA to 3xFLAG and introduced the gene encoding the fusion onto the chromosome at the *mbaA* locus. Tagging did not alter MbaA control over the biofilm lifecycle (Fig. S3A, Video 2). To inactivate the MbaA phosphodiesterase activity, we substituted the conserved catalytic residue E553 with alanine, converting EVL to AVL (referred to as EVL* henceforth). This change did not alter MbaA-3xFLAG abundance (Fig. S3B). *V. cholerae* harboring MbaA^EVL*^ exhibited an increase in biofilm biomass and a strong biofilm dispersal defect (Fig. 4A), and only a modest response to exogenous polyamines, with norspermidine eliciting some inhibition of biofilm dispersal and spermidine driving a small reduction in overall biofilm biomass (Fig. 4B, Video 2). Treatment with norspermidine did increase c-di-GMP levels in *V. cholerae* carrying MbaA^EVL*^ as judged by the reporter output (Fig. 4C), albeit not to the level of WT (Fig. 3C), suggesting that the *V. cholerae mbaA*^*EVL*\*^ mutant, which is incapable of c-di-GMP degradation, retains the capacity to synthesize some c-di-GMP via the MbaA SGDEF domain. In contrast, the *V. cholerae mbaA*^*EVL*\*^ strain displayed little reduction in c-di-GMP levels in response to spermidine, presumably because it lacks the phosphodiesterase activity required to degrade c-di-GMP (Fig. 4C). Thus, despite the fact that the purified cytoplasmic domain functioned only as a phosphodiesterase *in vitro*, our results suggest that MbaA is capable of synthesizing c-di-GMP in the presence of norspermidine. To validate this prediction, we altered the conserved catalytic residues D426 and E427 to alanine residues, yielding the inactive MbaA SGAAF variant (referred to as SGDEF*). These substitutions did not affect protein levels (Fig. S3B). In every regard, the *mbaA*^*SGDEF\**^ mutant behaved identically to the Δ*mbaA* mutant. The *V. cholerae mbaA*^*SGDEF\**^ mutant exhibited a modest biofilm dispersal defect (Fig. 4A) and biofilm biomass was reduced in response to exogenous norspermidine and spermidine (Fig. 4D, Video 2). Furthermore, addition of polyamines to the *mbaA*^*SGDEF\**^ mutant harboring the c-di-GMP reporter did not drive substantial changes to the reporter output (Fig. 4E). These results show that the MbaA SGDEF domain is indispensable for the MbaA response to polyamines.

Based on the results in Figs. 3 and 4, we conclude that the MbaA phosphodiesterase and diguanylate cyclase domains are both required for MbaA to respond properly to polyamines to regulate *V. cholerae* biofilm dispersal. We propose a model in which, elevated periplasmic norspermidine levels drive NspS to bind to MbaA. Consequently, MbaA phosphodiesterase activity is suppressed and the diguanylate cyclase activity dominates, which leads to c-di-GMP accumulation and commitment to the biofilm lifestyle (Fig. 4F). In contrast, when spermidine is detected, or when periplasmic polyamine concentrations are low, NspS dissociates from MbaA. Consequently, MbaA diguanylate cyclase activity is suppressed and phosphodiesterase activity dominates, which leads to a reduction in cytoplasmic c-di-GMP and biofilm dispersal (Fig. 4F).

**Fig. 4.**
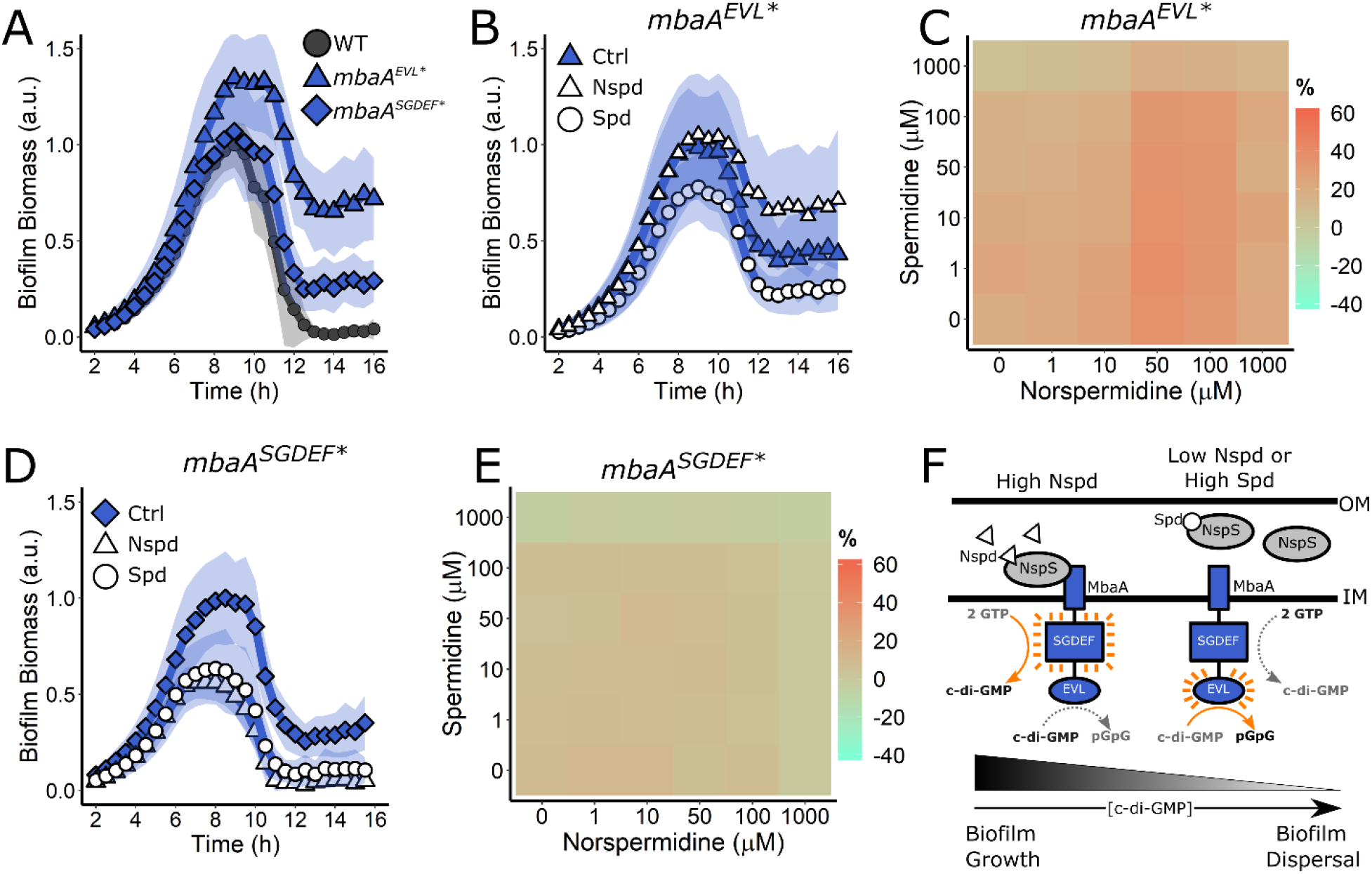
Both the MbaA SGDEF and EVL domains are required for regulation of *V. cholerae* biofilm dispersal. (A) Quantitation of biofilm biomass over time for the *V. cholerae* strains carrying *mbaA-3xFLAG, mbaA*^*EVL\**^*-3xFLAG*, and *mbaA*^*SGDEF\**^*-3xFLAG*. (B) Quantitation of biofilm biomass over time measured by time-lapse microscopy for *V. cholerae* carrying *mbaA*^*EVL\**^*-3xFLAG* following addition of a water (Ctrl), 100 µM norspermidine, or 100 µM spermidine. (C) C-di-GMP reporter output at the indicated polyamine concentrations for *V. cholerae* carrying *mbaA*^*EVL\**^*-3xFLAG*. Relative reporter signal (% difference) is displayed as a heatmap (teal and orange represent the lowest and highest reporter output, respectively). (D) As in B for the *mbaA*^*SGDEF\**^*-3xFLAG* mutant. (E) As in C for the *mbaA*^*SGDEF\**^*-3xFLAG* mutant. (F) Schematic representing the proposed MbaA activity in response to norspermidine and spermidine. Biofilm biomass data are represented as means normalized to the peak biofilm biomass of the WT strain or Ctrl condition. In all cases, *N* = 3 biological and *N* = 3 technical replicates, ± SD (shaded). a.u., arbitrary unit. In the c-di-GMP reporter assays, values are expressed as the percentage difference relative to the untreated WT strain, allowing comparisons to be made across all heatmaps in all figures in this manuscript. The same color bar applies to all panels in this manuscript. For each condition, *N* = 3 biological replicates. Numerical values and associated standard deviations are available in Table S1. In panel F, OM = outer membrane; IM = inner membrane.

### Polyamines control the *V. cholerae* biofilm lifecycle via periplasmic detection, not via import into the cytoplasm

Here, we investigate the role of PotD1 in controlling biofilm dispersal. Addition of norspermidine did not alter the non-dispersing phenotype of the Δ*potD1* mutant, whereas spermidine treatment drove premature biofilm dispersal and a reduction in peak biofilm biomass (Fig. 5A, Video 1). Consistent with these findings, addition of norspermidine to the Δ*potD1* strain did not alter the c-di-GMP reporter output, whereas addition of spermidine reduced output signal, but only at the highest concentrations tested (Fig. 5B). These biofilm lifecycle and reporter results show that PotD1-mediated import is not required for spermidine to modulate the biofilm lifecycle. Thus, we considered an alternative mechanism to underlie the *V. cholerae* Δ*potD1* phenotype. We hypothesized that endogenously produced norspermidine is secreted into the periplasm. In the case of WT *V. cholerae*, norspermidine is reimported by PotD1, leaving MbaA unliganded and predominantly in phosphodiesterase mode (Fig. 5C). In the *V. cholerae* Δ*potD1* mutant, we predict that periplasmic norspermidine levels become elevated, NpsS detects norspermidine, binds to MbaA, and promotes the MbaA diguanylate cyclase active state. Hence, c-di-GMP is synthesized (Fig. 5D) and biofilm dispersal is prevented. This hypothesis is consistent with our result showing that addition of norspermidine to the Δ*potD1* mutant does not cause any change in biofilm dispersal. Presumably, the NspS protein is already saturated due to elevated periplasmic levels of endogenously produced norspermidine. If our hypothesis is correct, then the Δ*mbaA* mutation should be epistatic to the Δ*potD1* mutation. Indeed, the Δ*mbaA* Δ*potD1* double mutant exhibited a dispersal phenotype indistinguishable from the single Δ*mbaA* mutant (Fig. 5E). Administration of exogenous norspermidine or spermidine to the Δ*mbaA* Δ*potD1* double mutant (Fig. 5F, Video 1) mimicked what occurred following addition to the single Δ*mbaA* mutant (Fig. 3D): that is, essentially no response.

**Fig. 5.**
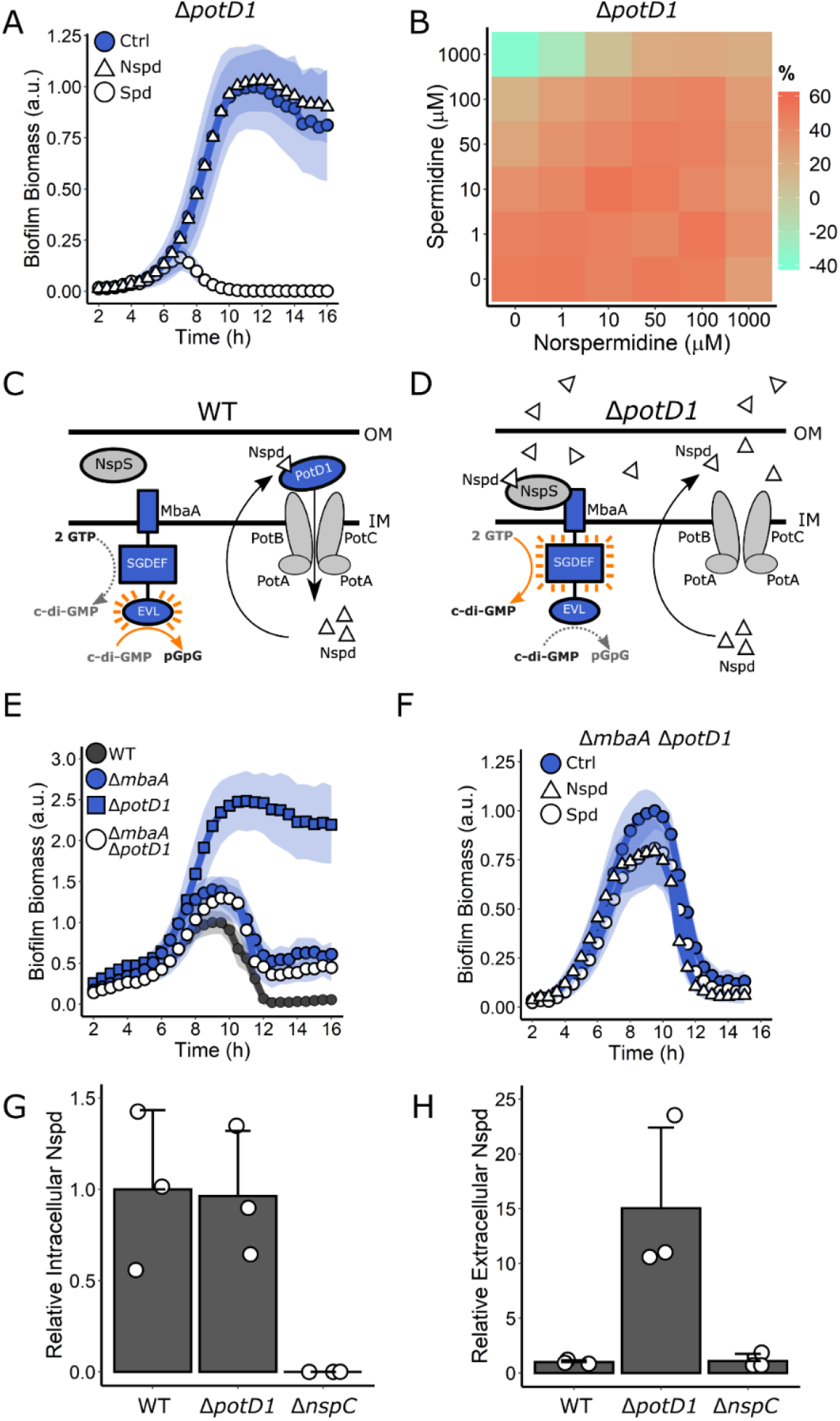
Polyamine import is not required for *V. cholerae* regulation of biofilm dispersal but is required to reduce external norspermidine levels. (A) Quantitation of biofilm biomass over time measured by time-lapse microscopy following addition of water (Ctrl), 100 µM norspermidine, or 100 µM spermidine to the Δ*potD1* mutant. (B) C-di-GMP reporter output at the indicated polyamine concentrations for the Δ*potD1* strain. Relative reporter signal (% difference) is displayed as a heatmap (teal and orange represent the lowest and highest reporter output, respectively). (C) Schematic of NspS-MbaA periplasmic detection of polyamines and polyamine import by PotD1 in WT *V. cholerae*. OM = outer membrane; IM = inner membrane. (D) Schematic of NspS-MbaA periplasmic detection of norspermidine and the accumulation of elevated extracellular norspermidine in the *V. cholerae* Δ*potD1* mutant. OM = outer membrane; IM = inner membrane. (E) Biofilm biomass over time for WT, the Δ*mbaA* mutant, the Δ*potD1* mutant, and the Δ*mbaA* Δ*potD1* double mutant. (F) As in A for the Δ*mbaA* Δ*potD1* double mutant. (G) Relative cytoplasmic norspermidine levels for WT, Δ*potD1*, and Δ*nspC* stains measured by mass spectrometry and normalized to the level present in the WT strain. (H) As in G for the corresponding cell-free culture fluids from the strains. The strains used in panels G and H also contained a Δ*vpsL* mutation to eliminate the possibility of trapping of norspermidine in the biofilm matrix. The data presented in panels G and H are internally normalized and comparisons should not be made across the two graphs. Biofilm biomass data are represented as means normalized to the peak biofilm biomass of the WT strain or Ctrl condition and in all cases, *N* = 3 biological and *N* = 3 technical replicates, ± SD (shaded). a.u., arbitrary unit. In the c-di-GMP reporter assays, values are expressed as the percentage difference relative to the untreated WT strain, allowing comparisons to be made across all heatmaps in all figures in the manuscript. The same color bar applies to all c-di-GMP reporter heatmaps in the manuscript and for each condition, *N* = 3 biological replicates. For c-di-GMP measurements, numerical values and associated standard deviations are available in Table S1. For norspermidine mass spectrometry measurements, *N* = 3 biological replicates, ± SD (all datapoints shown).

To garner additional evidence for the supposition that the Δ*potD1* strain possesses higher levels of extracellular norspermidine than does WT *V. cholerae*, we prepared cell-free culture fluids from the two strains and assessed norspermidine levels. We would expect that intracellular norspermidine levels should be equivalent in the two strains since reimport or lack thereof would not affect endogenous production. Thus, we also measured norspermidine levels in the companion cytoplasmic fractions. Mass spectrometry analyses showed that, indeed, the cytoplasmic norspermidine levels were identical in the WT and the Δ*potD1* strains (Fig. 5G). As a control we show that cytoplasmic norspermidine was nearly undetectable in a *V. cholerae* Δ*nspC* mutant that lacks the carboxynorspermidine decarboxylase enzyme, NspC, responsible for norspermidine synthesis (Fig. 5G) (Lee et al., 2009). In contrast, 15-fold more norspermidine could be extracted from cell-free culture fluids of the Δ*potD1* mutant than from WT *V. cholerae* (Fig. 5H). Thus, self-secreted norspermidine is internalized via PotD1. Together, these results demonstrate that polyamine signals exert their control over biofilm dispersal through the MbaA periplasmic sensing system and not through import nor some role in the cytoplasm.

## Discussion

In this study, we investigated the effects of the polyamines norspermidine and spermidine on *V. cholerae* biofilm dispersal. Norspermidine and spermidine, which differ in structure only by one methylene group, mediate starkly opposing effects on the *V. cholerae* biofilm lifecycle: norspermidine inhibits and spermidine promotes biofilm dispersal. Both function through the NspS-MbaA circuit. Thus, NspS, the polyamine binding protein must harbor the exquisite capability to assess the presence or absence of a single chemical moiety, as norspermidine binding drives NspS interaction with MbaA whereas binding to spermidine prevents this interaction. Here, we speculate on the possible biological significance of our findings. We suspect that norspermidine and spermidine act as ‘self’ and ‘other’ cues, respectively. Specifically, norspermidine is a rare polyamine in the biosphere, produced only by select organisms, namely *V. cholerae* and closely related marine vibrios (Hamana, 1997; Michael, 2018; Yamamoto et al., 1991). While WT *V. cholerae* releases little norspermidine, at least in part due to PotD1-mediated reuptake (Fig. 5C), it is possible that norspermidine is secreted by other vibrios, *V. cholerae* detects the released norspermidine via NpsS-MbaA, and its dispersal from biofilms is prevented. Thus, when close relatives are nearby, as judged by detection of the presence of norspermidine, *V. cholerae* elects to remain in its current biofilm niche. More speculative is the notion that norspermidine functions as a cue for phage-, predator-, or toxin-induced cell lysis. Specifically, if *V. cholerae* can detect norspermidine released by neighboring lysed *V. cholerae* cells, in response to the perceived danger, *V. cholerae* would remain in the protective biofilm state. Conversely, detection of spermidine, a nearly ubiquitously produced polyamine (Michael, 2018), could alert *V. cholerae* to the presence of competing or unrelated organisms. In this case, *V. cholerae* would respond by dispersing from biofilms and fleeing that locale. In a seemingly parallel scenario, we previously discovered that autoinducer AI-2, a universally produced quorum-sensing signal, also drove *V. cholerae* biofilm repression and premature dispersal, while CAI-1, the *V. cholerae* ‘kin’ quorum-sensing autoinducer did not (Bridges and Bassler, 2019). Together, our findings suggest that *V. cholerae* possesses two mechanisms to monitor its own numbers (norspermidine and CAI-1) and two mechanisms to take a census of unrelated organisms in the environment (spermidine and AI-2), and it responds by remaining in the biofilm state when closely related species are in the majority and by exiting the biofilm state when the density of non-related organisms is high, presumably to escape competition/danger.

We found that periplasmic polyamine detection by NspS-MbaA, but not polyamine import, drives the biofilm dispersal program and both the MbaA EVL and SGDEF domains are required for the *V. cholerae* response to polyamines. Despite previous work suggesting that MbaA functions exclusively as a phosphodiesterase and that its SGDEF domain, which possesses a serine substitution in the active site relative to the canonical GGDEF active site, is inert with respect to c-di-GMP synthesis, our data show that MbaA does synthesize c-di-GMP when norspermidine is present (Fig. 3C, Fig. 4C), as has also been shown for other SGDEF proteins (Pérez-Mendoza et al., 2011). Therefore, MbaA is a bifunctional protein capable of both c-di-GMP synthesis and degradation. We wonder how such an arrangement benefits *V. cholerae* in its response to polyamines. We speculate that the ability of MbaA to inversely alter c-di-GMP levels in response to two discrete cues coordinates polyamine signaling and enables more rapid and greater magnitude changes in c-di-GMP levels than would be possible if two separate receptors existed, each harboring only a single catalytic activity and each responsive to only one polyamine. The MbaA SGDEF and EVL domains could also participate in dimerization and/or allosteric binding to c-di-GMP or other metabolites to mediate feedback regulation of the opposing MbaA activities. Analogous examples exist, here we present one: *Caulobacter crescentus* contains a GGDEF/EAL protein called CC3396, in which the GGDEF domain is non-catalytic but it nonetheless binds to GTP, which allosterically activates the phosphodiesterase activity in the neighboring EAL domain (Christen et al., 2005). A feedback mechanism could provide MbaA with the ability to integrate the information encoded in extracellular polyamine blends with cytoplasmic cues, such as the metabolic state of the cell or the current cytoplasmic c-di-GMP level. Taken together, the double ligand detection capability linked to the dual catalytic activities of the NspS-MbaA circuit allows *V. cholerae* to distinguish between remarkably similar ligands and, in response, have the versatility to convey distinct information into the cell to alter the biofilm lifecycle.

In summary, four signaling pathways have now been defined that feed into the regulation of *V. cholerae* biofilm dispersal: starvation (RpoS) (Singh et al., 2017), quorum sensing (via CAI-1 and AI-2) (Bridges and Bassler, 2019; Singh et al., 2017), the recently identified DbfS/DbfR cascade (ligand unknown) (Bridges et al., 2020), and through the current work, polyamine signaling via NpsS-MbaA. We propose that integrating the information contained in these different stimuli into the control of biofilm dispersal endows *V. cholerae* with the ability to successfully evaluate multiple features of its fluctuating environment prior to committing to the launch of this key lifestyle transition that, ultimately, impinges on its overall survival. Because *V. cholerae* biofilm formation and biofilm dispersal are intimately connected with cholera disease and its transmission, we suggest that deliberately controlling the biofilm lifecycle, possibly via synthetic strategies that target polyamine signal transduction via NspS-MbaA could be a viable therapeutic strategy to ameliorate disease.

## Materials and Methods

### Bacterial strains, reagents, and imaging assays

The *V. cholerae* strain used in this study was WT O1 El Tor biotype C6706str2. Antibiotics were used at the following concentrations: polymyxin B, 50 μg/mL; kanamycin, 50 μg/mL; spectinomycin, 200 μg/mL; and chloramphenicol, 1 μg/mL. Strains were propagated for cloning purposes in lysogeny broth (LB) supplemented with 1.5% agar or in liquid LB with shaking at 30°C. For biofilm dispersal analyses, *lux* measurements, c-di-GMP reporter quantitation, and norspermidine measurements, *V. cholerae* strains were grown in M9 medium supplemented with 0.5% dextrose and 0.5% casamino acids. All strains used in this work are reported in Table S2. Compounds were added from the onset of biofilm initiation. Norspermidine (Millipore Sigma, I1006-100G-A) and spermidine (Millipore Sigma, S2626-1G) were added at the final concentrations designated in the figures. The biofilm lifecycle was measured using time-lapse microscopy as described previously (Bridges and Bassler, 2019). All plots were generated using ggplot2 in R. Light production driven by the *vpsL* promoter was monitored as described previously (Bridges et al., 2020). Results from replicates were averaged and plotted using ggplot2 in R.

### DNA manipulation and strain construction

All strains generated in this work were constructed by replacing genomic DNA with DNA introduced by natural transformation as previously described (Bridges et al., 2020). PCR and Sanger sequencing were used to verify correct integration events. Genomic DNA from recombinant strains was used for future co-transformations and as templates for PCR to generate DNA fragments, when necessary. See Table S3 for primers and g-blocks (IDT) used in this study. Gene deletions were constructed in frame and eliminated the entire coding sequences. The exceptions were *mbaA* and *nspS*, which overlap with adjacent genes, so an internal portion of each gene was deleted, ensuring that adjacent genes were not perturbed. A similar strategy was used for 3x-FLAG tagging of MbaA (see Table S3 for details). All strains constructed in this study were verified using the Genewiz sequencing service.

### Western blotting

Cultures of strains carrying MbaA-3xFLAG (and relevant catalytic site variants) were collected at OD_600_ = 1.0 and subjected to centrifugation for 1 min at 13,000 rpm. The pellets were flash frozen, thawed for 5 min at 25^°^ C, and subsequently chemically lysed by resuspending to OD_600_ = 1.0 in 75 μL Bug Buster (Novagen, #70584–4) supplemented with 0.5% Triton-X, 50 μg/mL lysozyme, 25 U/mL benzonase nuclease, and 1 mM phenylmethylsulfonyl fluoride (PMSF) for 10 min at 25^°^ C. Lysates were solubilized in 1X SDS-PAGE buffer for 1 h at 37^°^ C. Samples were loaded into 4–20% Mini-Protein TGX gels (Bio-Rad). Electrophoresis was carried out at 200 V. Proteins were transferred from the gels to PVDF membranes (Bio-Rad) for 50 min at 4^°^ C at 100 V in 25 mM Tris base, 190 mM glycine, 20% methanol. Following transfer, membranes were blocked in 5% milk in PBST (137 mM NaCl, 2.7 mM KCl, 8 mM Na_2_HPO_4_, 2 mM KH_2_PO_4_, and 0.1% Tween) for 1 h, followed by three washes with PBST. Subsequently, membranes were incubated for 1 h with a monoclonal Anti-FLAG-Peroxidase antibody (Millipore Sigma, #A8592) at a 1:5,000 dilution in PBST with 5% milk. After washing six times with PBST for 5 min each, membranes were exposed using the Amersham ECL Western blotting detection reagent (GE Healthcare). For the RpoA loading control, the same protocol was followed except that the primary antibody was Anti-*Escherichia coli* RNA Polymerase α (Biolegend, #663104) used at a 1:10,000 dilution and the secondary antibody was an Anti-Mouse IgG HRP conjugate antibody (Promega, #W4021) used at a 1:10,000 dilution.

### Extraction and measurement of norspermidine

To measure intracellular norspermidine, previously established techniques were followed with slight modifications (McGinnis et al., 2009). The indicated *V. cholerae* strains were grown in M9 medium containing glucose and casamino acids. 0.5 OD_600_ equivalents of *V. cholerae* cells were collected by centrifugation for 1 min at 13,000 rpm from cultures that had been grown to OD_600_ ∼2.0. Cells were washed once with 1x PBS and subsequently resuspended in 50 µL PBS. The resuspended cells were lysed by 10x freeze-thaw cycles in liquid nitrogen followed by 20 sec of bath sonication (Fisher Scientific, FS30). To precipitate proteins, 15 µL of 40% trichloroacetic acid was added to the lysates and the mixtures were incubated for 5 min. Precipitated material was pelleted by centrifugation for 5 min at 13,000 rpm. 50 µL of the resulting clarified supernatants were transferred to new microcentrifuge tubes and frozen at −20^°^ C until further analysis. For measurements of norspermidine in cell-free culture fluids, *V. cholerae* strains were grown as a above and culture fluids were collected following the centrifugation step. Remaining cells were removed by filtration with 0.45 µm filters (Millex-HV), and proteins were precipitated as above. In all cases, synthetic norspermidine was simultaneously analyzed as a control.

Samples were derivatized with benzoyl chloride as previously described (Morgan, 1998). HPLC was performed on a Shimadzu UFLC system with PAL autoinjector. Thirty minute-gradient chromatography separation was performed using solvent A (10% methanol / 90% water) and solvent B (90% methanol / 10% water) on an ACE Ultracore 2.5 SuperC18 1.0 × 50 mm column with 45 µL/min flow rate at a column temperature of 45^°^ C. Mass spectrometry was performed using an Orbitrap XL mass spectrometer (Thermo) with an APCI ionization source in positive mode. The parent ion (MS1) was detected in the Orbitrap with 100,000 mass resolution and fragment ions (MS2) were detected in the ion trap. Parameters were as follows: the vaporizer temperature was 270^°^ C, the sheath gas flow rate was 18 a.u., the auxiliary gas flow rate was 5 a.u., the sweep gas flow rate was 5 a.u., the discharge current was 9 mA, the capillary temperature was 250^°^ C, the capillary voltage was 36 V, and the tube lens was 72 V. Skyline software (University of Washington) was used to analyze results and values were normalized to the OD_600_ of the culture when it was collected.

### c-di-GMP reporter assays

The c-di-GMP reporter has been described (Zamorano-Sánchez et al., 2019; Zhou et al., 2016). To measure relative reporter output for each condition, 100 µL of *V. cholerae* cultures were back diluted to OD_600_ = 0.0002 following overnight growth. Cultures were dispensed into 96-well plates containing the indicated polyamines and the plates were covered in breathe-easy membranes to prevent evaporation. Samples were incubated overnight at 30^°^ C with shaking. The following morning, the breathe-easy membranes were removed and fluorescence measurements were obtained using a BioTek Synergy Neo2 Multi-Mode reader. For AmCyan, the excitation wavelength was 440±20 nm and emission was detected at 490±20 nm and for TurboRFP, the excitation wavelength was 530±20 nm and the emission was 575±20 nm. The c-di-GMP-regulated TurboRFP fluorescence was divided by the constitutive AmCyan fluorescence to yield the relative fluorescence intensity (RFI). To facilitate comparisons between strains and conditions, RFIs were subsequently normalized to the untreated *V. cholerae* WT RFI and the data are expressed as the percentage differences (denoted relative reporter signal). All results were obtained in biological triplicate and data analysis and plotting were performed in R.

## Acknowledgements

We thank members of the Bassler group and Prof. Ned Wingreen for thoughtful discussions. The c-di-GMP reporter plasmid was a gift from Fitnat Yildiz (UC Santa Cruz). Mass spectrometry was conducted by the Princeton Molecular Biology Proteomics and Mass Spectrometry Core Facility. This work was supported by the Howard Hughes Medical Institute, NIH Grant 5R37GM065859, National Science Foundation Grant MCB-1713731, and a Max Planck-Alexander von Humboldt research award to BLB. AAB is a Howard Hughes Medical Institute Fellow of the Damon Runyon Cancer Research Foundation, DRG-2302-17. The content is solely the responsibility of the authors and does not necessarily represent the official views of the National Institutes of Health. The funders had no role in study design, data collection and analysis, decision to publish, or preparation of the manuscript.

---

**Fig. S1.**
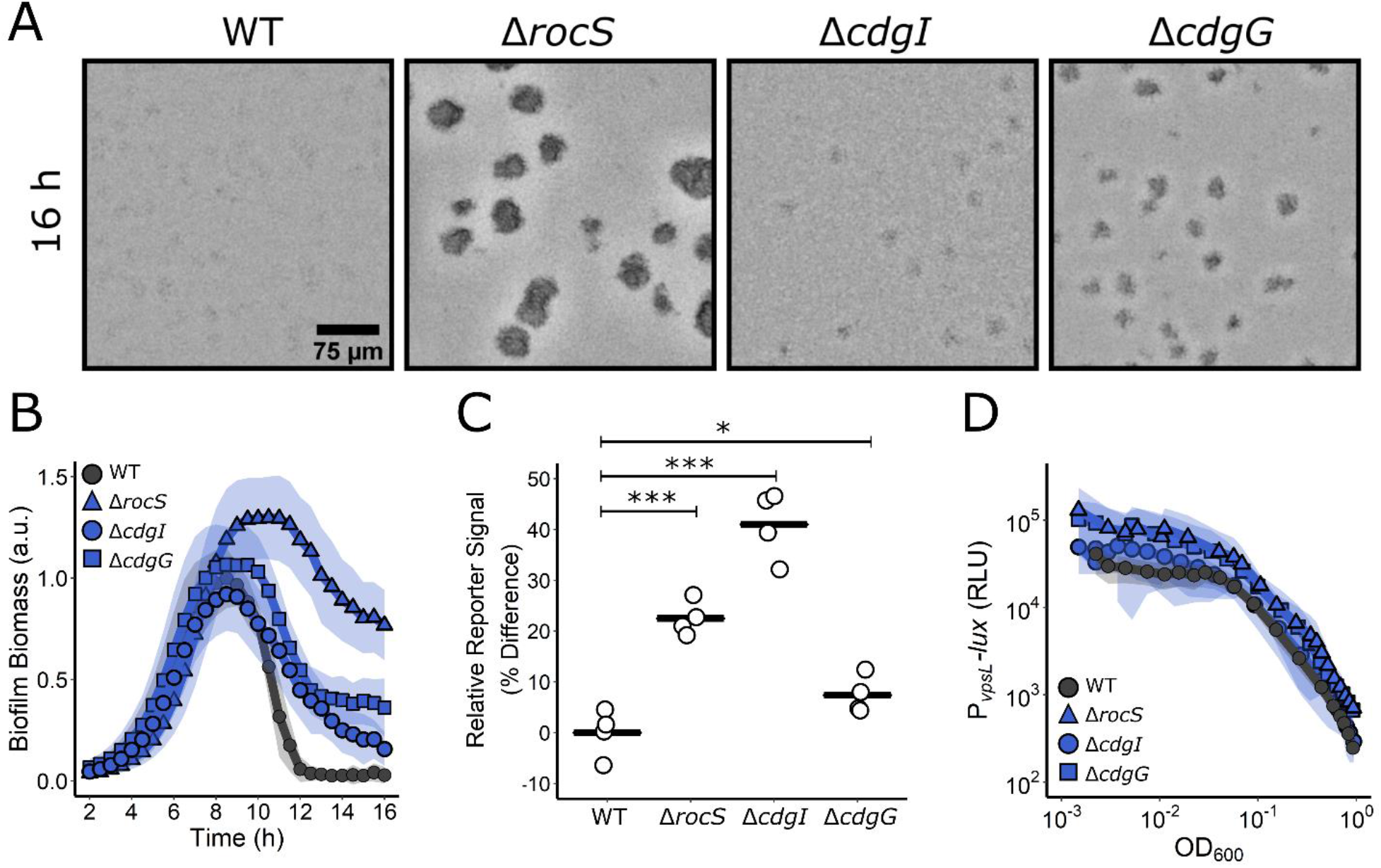
C-di-GMP signaling controls biofilm dispersal. (A) Representative brightfield images of biofilms at 16 h for the designated strains. (B) Quantitation of biofilm biomass over time measured by time-lapse microscopy for WT and the designated mutants. Data are represented as means normalized to the peak biofilm biomass of the WT strain. *N* = 3 biological and *N* = 3 technical replicates, ± SD (shaded). a.u., arbitrary unit. (C) Relative c-di-GMP reporter signal for the indicated strains. Values are expressed as the percentage difference relative to the WT strain. *N* = 4 biological replicates. The black bar shows the sample mean. Unpaired t-tests were performed for statistical analysis, with P values denoted as *P < 0.05; *** P < 0.0005. (D) The corresponding *PvpsL-lux* reporter output for the strains and growth conditions in B over the growth curve. RLU, relative light units. *N* = 3 biological replicates, ± SD (shaded).

**Fig. S2.**
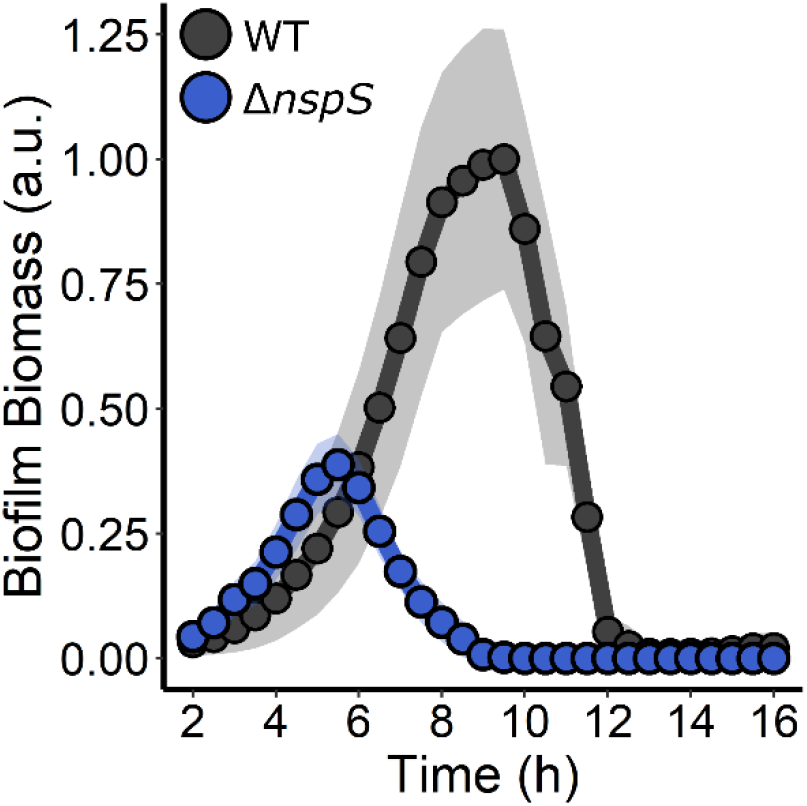
NspS is required for *V. cholerae* biofilm formation. Quantitation of biofilm biomass over time measured by time-lapse microscopy for WT *V. cholerae* and the Δ*nspS* strain. For each strain, *N* = 3 biological and *N* = 3 technical replicates, ± SD (shaded). a.u., arbitrary unit.

**Fig. S3.**
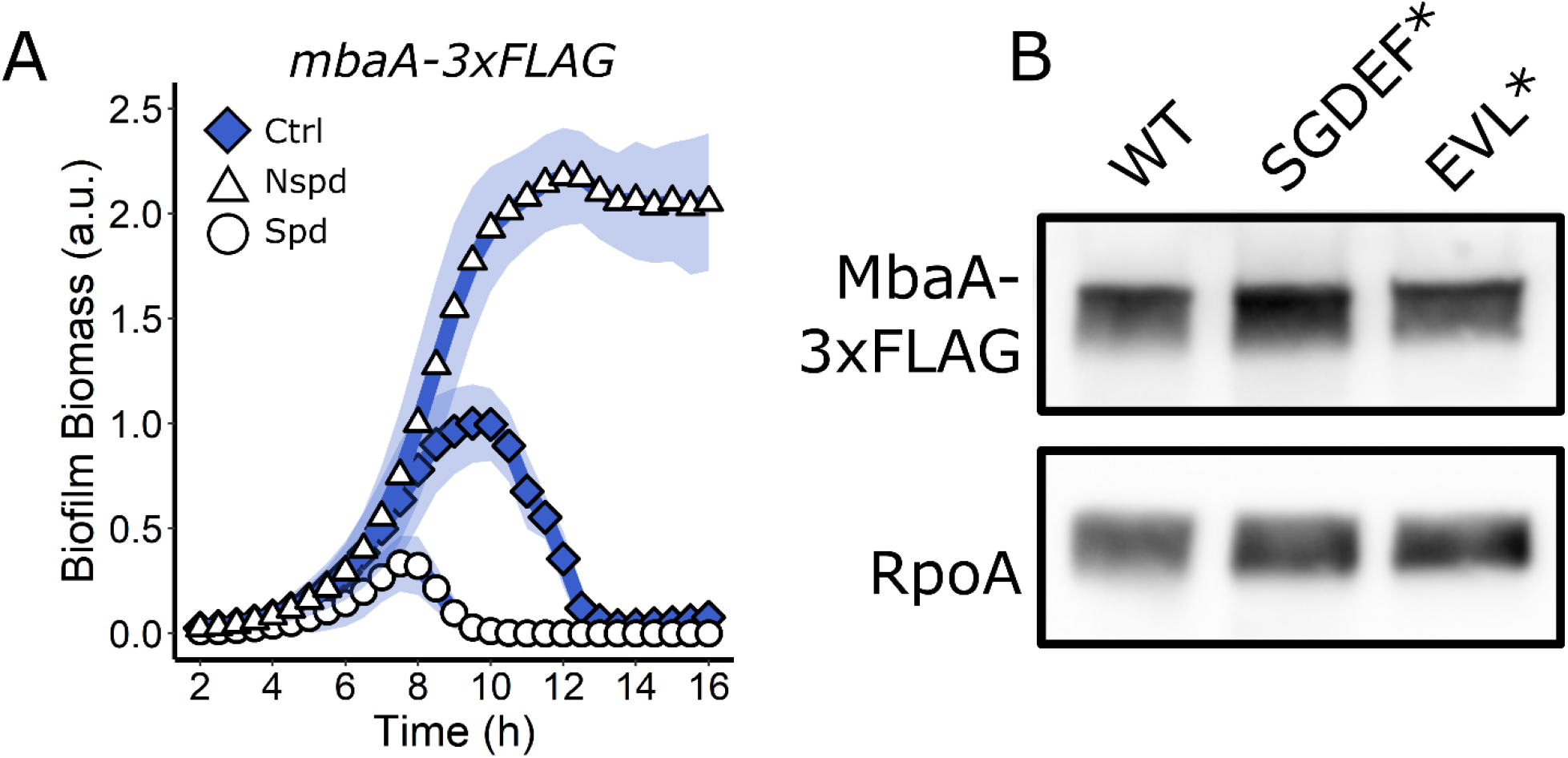
MbaA active site mutations do not alter protein levels. (A) Quantitation of biofilm biomass over time measured by time-lapse microscopy for *V. cholerae* carrying MbaA-3xFLAG following addition of a water (Ctrl), 100 µM norspermidine, or 100 µM spermidine. Biofilm biomass data are represented as means normalized to the peak biofilm biomass of the WT strain or Ctrl condition. In all cases, *N* = 3 biological and *N* = 3 technical replicates, ± SD (shaded). a.u., arbitrary unit. (B) Top panel: Western blot showing 3xFLAG for the WT MbaA, MbaA SGDEF*, and MbaA EVL* proteins. Bottom panel: RpoA is the loading control. Data are representative of three biological replicates.

**Table S1. (separate file). C-di-GMP reporter output averages and standard deviations.**

**Table S2.** Strains used in this study.

| Strain Number | Genotype | Plasmid | Antibiotic Resistance | Origin |
| --- | --- | --- | --- | --- |
| BB_Vc_0090 | WT O1 El Tor biotype C6706str2 | - | Sm | Laboratory WT |
| AB_Vc_761 | $\Delta vc1807::Cm^R$ (Referred to as WT) | - | Sm, Cm | Bridges et al. 2020 |
| AB_Vc_839 | $\Delta mbaA \Delta vc1807::Kan^R$ | - | Sm, Kan | Bridges et al. 2020 |
| AB_Vc_928 | $\Delta nspS \Delta vc1807::Kan^R$ | | Sm, Kan | NT of BB_Vc_0090 |
| AB_Vc_711 | $\Delta potD1 \Delta vc1807::Cm^R$ | - | Sm, Cm | Bridges et al. 2020 |
| AB_Vc_758 | $\Delta rocS \Delta vc1807::Cm^R$ | - | Sm, Cm | Bridges et al. 2020 |
| AB_Vc_777 | $\Delta cdgI \Delta vc1807::Cm^R$ | - | Sm, Cm | Bridges et al. 2020 |
| AB_Vc_778 | $\Delta cdgG \Delta vc1807::Cm^R$ | - | Sm, Cm | Bridges et al. 2020 |
| AB_Vc_835 | $mbaA-3xFLAG \Delta vc1807::Kan^R$ | - | Sm, Cm | NT of BB_Vc_0090 |
| AB_Vc_870 | $mbaA(D426A, E427A)-3xFLAG \Delta vc1807::Kan^R$ | - | Sm, Cm | NT of BB_Vc_0090 |
| AB_Vc_872 | $mbaA(E553A)-3xFLAG \Delta vc1807::Kan^R$ | - | Sm, Cm | NT of BB_Vc_0090 |
| AB_Vc_798 | $\Delta mbaA \Delta potD1 \Delta vc1807::Kan^R$ | - | Sm, Kan | NT of AB_Vc_709 |
| AB_Vc_956 | WT | pFY4357::Gm<sup>R</sup> | Sm, Gm | Conj. of BB_Vc_0090 |
| AB_Vc_975 | $\Delta mbaA \Delta vc1807::Kan^R$ | pFY4357::Gm<sup>R</sup> | Sm, Gm, Kan | Conj of AB_Vc_839 |
| AB_Vc_977 | $\Delta nspS \Delta vc1807::Kan^R$ | pFY4357::Gm<sup>R</sup> | Sm, Gm, Kan | Conj. of AB_Vc_928 |
| AB_Vc_971 | $mbaA(D426A, E427A)-3xFLAG \Delta vc1807::Kan^R$ | pFY4357::Gm<sup>R</sup> | Sm, Gm, Kan | Conj. of AB_Vc_870 |
| AB_Vc_973 | $mbaA(E553A)-3xFLAG \Delta vc1807::Kan^R$ | pFY4357::Gm<sup>R</sup> | Sm, Gm, Kan | Conj. of AB_Vc_872 |
| AB_Vc_969 | $\Delta potD1 \Delta vc1807::Cm^R$ | pFY4357::Gm<sup>R</sup> | Sm, Gm, Kan | Conj. of AB_Vc_711 |
| AB_Vc_987 | $\Delta rocS \Delta vc1807::Cm^R$ | pFY4357::Gm<sup>R</sup> | Sm, Gm, Kan | Conj. of AB_Vc_758 |
| AB_Vc_989 | $\Delta cdgI \Delta vc1807::Cm^R$ | pFY4357::Gm<sup>R</sup> | Sm, Gm, Kan | Conj. of AB_Vc_777 |
| AB_Vc_991 | $\Delta cdgG \Delta vc1807::Cm^R$ | pFY4357::Gm<sup>R</sup> | Sm, Gm, Kan | Conj. of AB_Vc_778 |
| AB_Vc_801 | $\Delta vc1807::Kan^R$ | pEVS143-*P<sub>vpsL</sub>*-*lux*::Cm<sup>R</sup> | Sm, Cm, Kan | Bridges et al. 2020 |
| AB_Vc_848 | $\Delta mbaA \Delta vc1807::Kan^R$ | pEVS143-*P<sub>vpsL</sub>*-*lux*::Cm<sup>R</sup> | Sm, Cm, Kan | Conj. of AB_Vc_839 |
| AB_Vc_828 | $\Delta potD1 \Delta vc1807::Kan^R$ | pEVS143-*P<sub>vpsL</sub>*-*lux*::Cm<sup>R</sup> | Sm, Cm, Kan | Conj. of AB_Vc_810 |
| AB_Vc_830 | $\Delta rocS \Delta vc1807::Kan^R$ | pEVS143-*P<sub>vpsL</sub>*-*lux*::Cm<sup>R</sup> | Sm, Cm, Kan | Conj. of AB_Vc_812 |
| AB_Vc_831 | $\Delta cdgI \Delta vc1807::Kan^R$ | pEVS143-*P<sub>vpsL</sub>*-*lux*::Cm<sup>R</sup> | Sm, Cm, Kan | Conj. of AB_Vc_813 |
| AB_Vc_832 | $\Delta cdgG \Delta vc1807::Kan^R$ | pEVS143-*P<sub>vpsL</sub>*-*lux*::Cm<sup>R</sup> | Sm, Cm, Kan | Conj. of AB_Vc_814 |
| AB_Vc_979 | $\Delta vpsL::Cm^R$ | - | Sm, Cm | NT of BB_Vc_0090 |
| AB_Vc_981 | $\Delta vpsL::Cm^R \Delta potD1 \Delta vc1807::Kan^R$ | - | Sm, Cm, Kan | NT of AB_Vc_810 |
| AB_Vc_983 | $\Delta vpsL::Cm^R \Delta nspC \Delta vc1807::Kan^R$ | - | Sm, Cm, Kan | NT of AB_Vc_823 |

NT = Natural Transformation
Conj = Conjugation

**Table S3.**
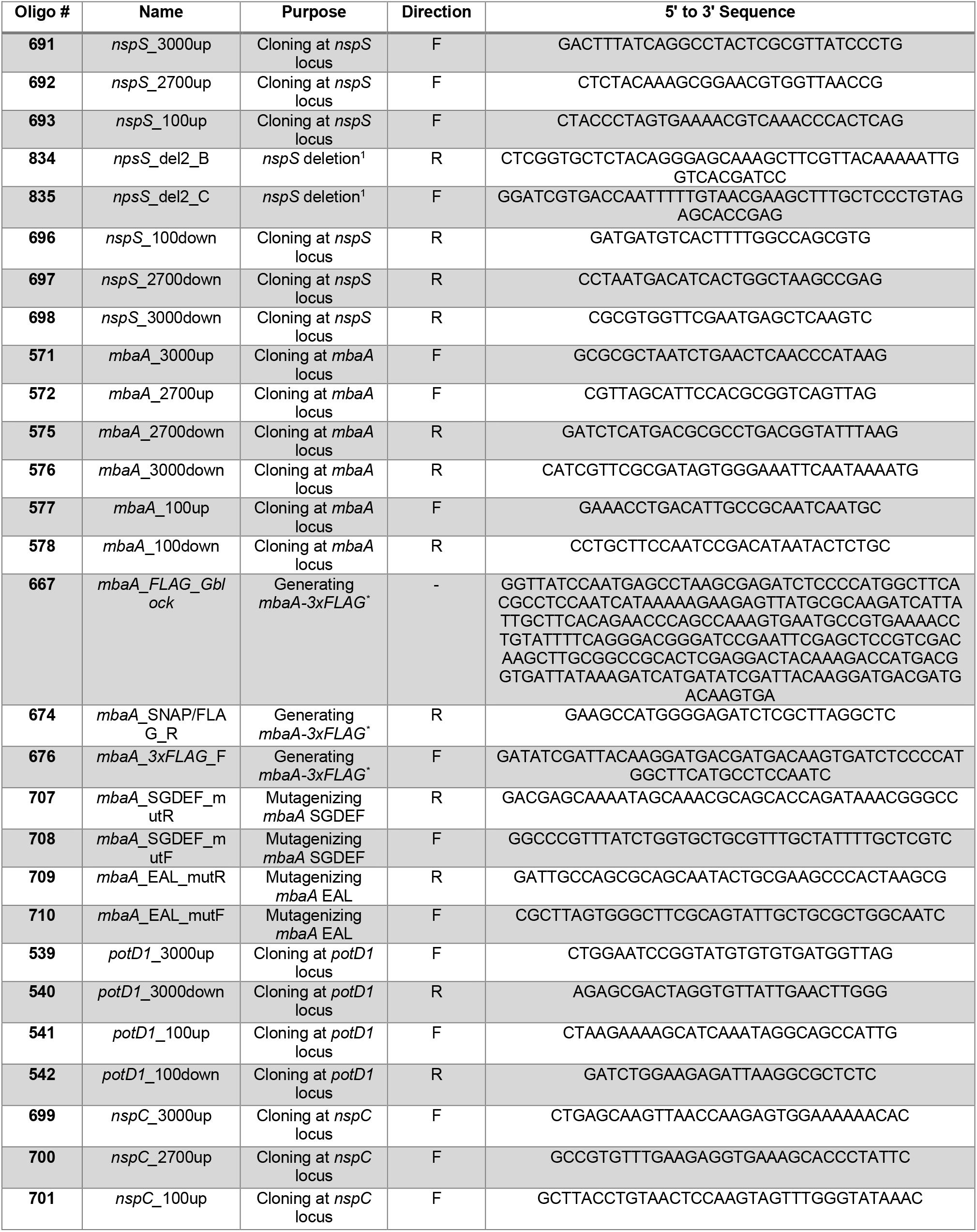
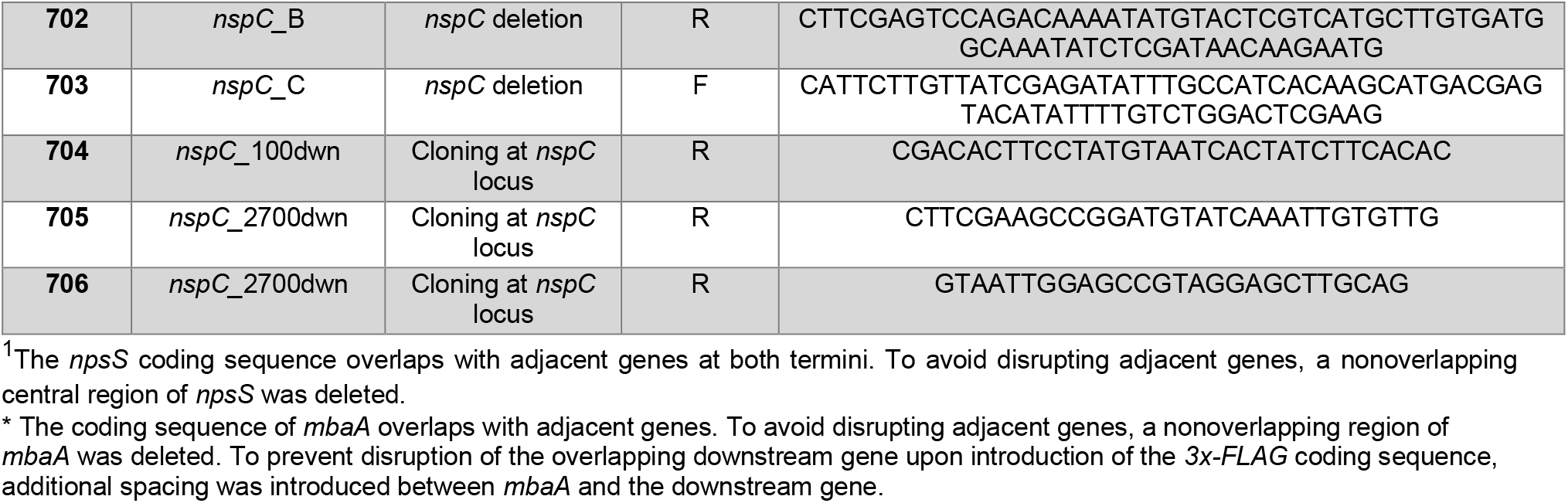
DNA oligonucleotides and gene fragments used in this study.

**Video 1 (separate file)**. Representative time-lapse images of the biofilm lifecycles of the WT, Δ*mbaA*, Δ*potD1*, and Δ*mbaA* Δ*potD1 V. cholerae* strains following treatment with water (Ctrl), 100 µM norspermidine (Nspd), or 100 µM spermidine (Spd).

**Video 2 (separate file)**. Representative time-lapse images of the biofilm lifecycles of the *mbaA-3xFLAG, mbaA*^*EVL\**^*-3xFLAG, mbaA*^*SGDEF\**^*-3xFLAG V. cholerae* strains following treatment with water (Ctrl), 100 µM norspermidine (Nspd), or 100 µM spermidine (Spd).

